# Differing ecological responses of seabirds to invasive species eradication

**DOI:** 10.1101/2021.04.07.438878

**Authors:** Jeremy P. Bird, Richard A. Fuller, Justine D. S. Shaw

## Abstract

The impact of invasive species at seabird breeding islands causes a breakdown of important ecological functions such as prey consumption and nutrient transfer, and elevates extinction risk in impacted taxa. Eradicating invasive species from islands can result in substantial short-term recovery of seabird populations and consequently the prevalence of eradication programs as conservation tools is increasing. However, as the scale and complexity of eradications has increased, quantitative data on rates of recovery, especially from larger islands, remain limited. Furthermore, the mechanisms that govern recovery are poorly understood, limiting our ability to forecast outcomes and therefore prioritise effectively. Here, using the world’s largest multi-species vertebrate eradication from Macquarie Island as a case study, we show how responses to invasive species and their eradication differ. Species with broad realised niches whose breeding phenology minimizes time on land and corresponds with summer resource abundance remained extant alongside invasive species while more habitat-specific species present in winter were extirpated. Following eradication, immigration and flexibility to colonise under-utilised optimal habitat appears to be boosting population growth in recolonising species, whereas established populations appear to be tethered to refugial habitats by the influence of philopatry, and their recovery is slower as a result. Unpicking these differential responses and the mechanisms behind them provides valuable information to help predict responses in other systems as future eradications are planned.

## MAIN

In a review of seabird responses to invasive species removal from islands Brooke et al observed more rapid population growth on islands where species were newly colonising than on islands where an established source population remained extant. They hypothesised that immigration plays a greater role than has been credited in seabird population recovery. We tested this theory in a natural experiment comparing the recovery of two established species and two recolonising species following the world’s largest multi-species eradication project on Macquarie Island.

Invasive species are a global problem. They are one of the most severe drivers of biodiversity loss, precipitating more species extinctions to date than other threats, and their impacts extend to fisheries, agriculture and human health (Reaser et al., 2007). Their impacts are particularly acute on islands, which hold a disproportionate amount of total, and threatened biodiversity (Russell and Kueffer, 2019). Huge progress has been made to develop the whole-island eradication of invasive species as a conservation tool, which has saved more species from extinction than any other management intervention (Bolam et al., 2020). However, in two ways, we are still at the start of the ecosystem recovery curve. First, the scale of the problem globally remains immense. Having honed our skills on uninhabited, relatively small and ecologically simple islands, we must now evolve to tackle ever larger, more complex, and human inhabited islands (Keitt et al., 2018). With such islands comes greater uncertainty around what the ecological outcomes will be. Second, full recovery may take decades. There is a pervasive assumption that, rid of invasive vertebrates, islands will recover to their pre-invaded state. This is not always the case (Prior et al., 2018), and further conservation interventions might be needed after the eradications themselves. Threatened species can become secure rapidly following an eradication, but recovery to abundant, pre-invasion maxima, and the restoration of their ecological functions might require much longer timeframes and additional management efforts. In all but a few cases, there has been insufficient time to determine how recovery is unfolding, and what post-eradication stable states islands can achieve.

In this context, every eradication is a learning opportunity, but systematic multi-taxa monitoring of post-eradication responses is rare (Towns, 2018). Both time to recovery and uncertainty of outcomes scale with island size and ecological complexity (Keitt et al., 2018). There is a need, therefore, to understand proximal outcomes, and the mechanisms that govern them, to be able to predict long-term outcomes. By reducing uncertainty, we can improve prioritisation and systematic conservation planning of future eradications. Here we utilise a rare opportunity of having time series data for multiple species to compare responses of a threatened seabird fauna to staged eradications of all vertebrate pests from Macquarie Island, the largest such intervention to date.

Procellariiform petrels are a particularly important group in island restoration. They are largely restricted to islands, where they have become disproportionately threatened with extinction, but under natural circumstances are influential ecosystem engineers. Highly susceptible to invasive species impacts 52 of 124 species (42%) are listed as threatened with extinction globally (Rodríguez et al., 2019). Paradoxically some petrels are still the most populous seabirds, consuming prey volumes commensurate with global fisheries, and transferring marine nutrients from pelagic to terrestrial and coastal systems in volumes equivalent to other major geochemical fluxes (Danckwerts et al., 2014; Otero et al., 2018).

Research following island eradication efforts shows that responses by petrels can be difficult to predict, suggesting that the ecological mechanisms at play are not fully understood (Buxton et al., 2014). For example, having been extirpated from Macquarie Island, Grey Petrels recolonised immediately as cat numbers were reduced through management and increased rapidly in the early years following eradication (Robinson and Copson, 2014; Schulz et al., 2006). Conversely, the population on Campbell Island, which lies closer to a source population at the Antipodes, remained small 14 years after rat eradication and had not recovered to the extent seen on Macquarie (Parker et al., 2017). It remains uncertain why responses apparently differed between the two islands, for example whether population change is driven purely by intrinsic growth or if immigration is also fuelling recovery. Such uncertainty impedes decision-making around future eradications and indicates a strong need to monitor the variability in responses between species to understand the ecological mechanisms at play.

Seabirds are naïve to invasive alien predators. Vulnerability scales loosely with size. Storm-petrels, the smallest species, rarely persist in the presence of even the smallest rodents, mice (Bolton et al., 2014), although there are an increasing number of examples of mice adapting to predate larger species too (e.g. Dilley et al., 2018). Conversely, larger predators such as cats are unable to enter the burrows of small-bodied species. Predators are often food limited in winter when summer breeding species are away from breeding grounds (Barbraud et al., 2009) so breeding phenology also affects susceptibility to invasive species. Different mechanisms govern recovery. Although seabirds are generally philopatric Brooke et al. (2018) have shown the importance of immigration in post-eradication population growth. While the influence of immigration in driving population growth is easy to detect—if growth is above the intrinsic rate of population growth immigration must be happening—the role of philopatry in slowing population growth by tethering colonies to sub-optimal habitat has not been considered.

In this paper we identify the interacting roles of body size, breeding phenology, immigration and philopatry in shaping the populations of four petrel species by influencing (i) susceptibility to invasive species impacts, (ii) timing and magnitude of responses to eradication, and (iii) their current distributions and habitat use.

## Results and discussion

We completed whole-island surveys of two established species that remained extant in the presence of invasive predators, Antarctic Prions *Pachyptila desolata* and White-headed Petrels *Pterodroma lessonii*, and two recolonising species extirpated in the 1900s, Blue Petrels *Halobaena caerulea* and Grey Petrels *Procellaria cineria*. We mapped their contemporary distribution and abundance (Bird et al. submitted A) and compared this with the only previous island-wide study from the late 1970s (Brothers, 1984). We also analysed population trends in monitoring sites first surveyed between 1963 and the mid-2000s (Brothers and Bone, 2008; Schulz et al., 2006).

All four species are now increasing (Figure 1). With the exception of Grey Petrels, current population sizes (Table 1), combined with positive trends sustained for >5 years suggests that Blue and White-headed Petrels are no longer threatened and based on established criteria both can now be delisted from Tasmania’s Threatened Species Protection Act 1995 (TSPA) and Australia’s federal Environment Protection and Biodiversity Conservation Act 1999 (EPBC). With a population of <1,000 mature individuals, Grey Petrels still meet the criteria for Vulnerable in Australia (irrespective of their increasing population trend). However, based upon current rates of population growth, and a population of 252 breeding pairs (~500 mature individuals) in 2018, we project they will exceed this threshold by 2026, possibly sooner if breeding occurs in the high proportion of non-breeding burrows recorded during surveys (Bird et al. submitted A).

**Figure 1:**
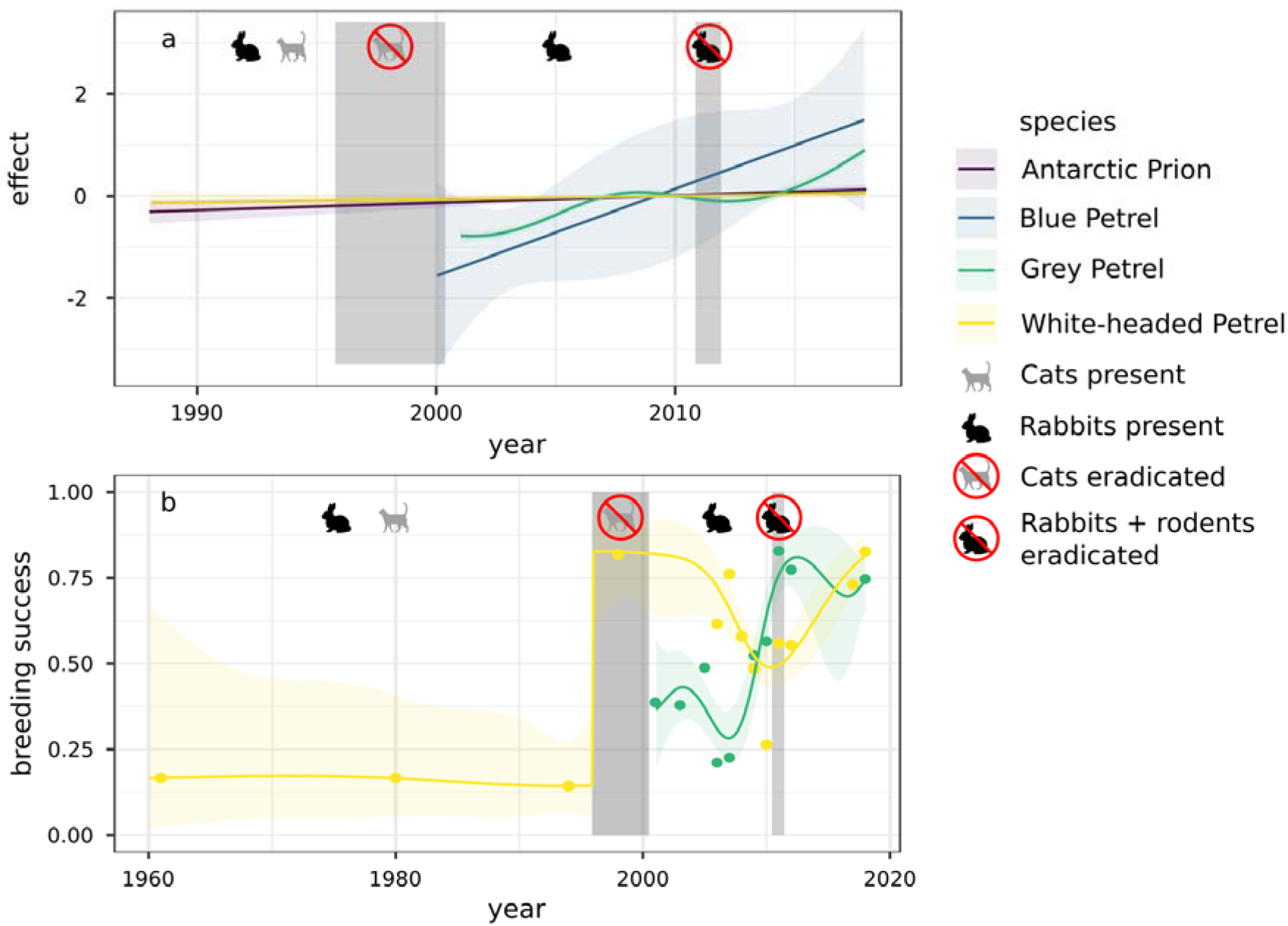
Fitted smooths of a) overall burrow numbers, and b) breeding success of petrels on Macquarie Island. The breeding success model included cats/no cats as a factor which allowed for the step change.

**Table 1:** Burrow numbers, breeding pairs ± 95% CIs (†Bird et al. submitted A), estimated occupancy (*Bird et al. submitted B) and population growth rate.

| Species | Current burrow estimate† | Occupancy* | Current breeding pairs† | Pop. growth |
| --- | --- | --- | --- | --- |
| | | | | rate ($\lambda$) |
| White-headed Petrel | 27,965 (20,223-38,669) | 0.46 | 12,769 (9,017-18,083) | 1.01 |
| Antarctic Prion | 239,362 (211,524-270,864) | 0.67 | 159,575 (140,710-180,968) | 1.01 |
| Blue Petrel | 11,898 (10,981-12,816) | 0.47 | 5,588 (5,042-6,326) | 1.14 |
| Grey Petrel | 630 (-) | 0.40 | 252 (227-302) | 1.10 |

This is a major outcome that marks the achievement of a proximal goal of pest management at Macquarie: to avert further seabird extinctions and promote threatened species recovery (Parks and Wildlife Service, 2014). It is important that this is communicated widely, and threatened species are downlisted. There is often emphasis on reporting these management outcomes which carry financial and policy implications, but less focus on ecological outcomes. In our study system comparison between species responses provides deeper ecological insights which we address below.

### i) Susceptibility to invasive species impacts

Species’ unique ecologies have shaped their responses to invasive species impacts on Macquarie Island. Throughout the period of invasive species occurrence Antarctic Prions and White-headed Petrels remained extant, whereas Blue and Grey Petrels were both extirpated in the 1900s (Brothers and Bone, 2008; Brothers, 1984).

Antarctic Prions were impacted the least. In a year, they spend the shortest amount of time at the island of any of the four species and they breed in the summer months when predators are not food limited (Brothers, 1984). Despite high adult mortality from cats and an unquantified level of nest predation from rats breeding success was apparently high (Brothers, 1984). Prion burrows are too small to be entered by cats, affording protection, and high adult mortality may be skewed towards non-breeding birds which, in other species, spend a disproportionate amount of time outside burrow entrances (Bonnaud et al., 2009). Brothers (1984) estimated prions were the most abundant species in the 1970s, with 48,900 breeding pairs, and they remain so today (Table 1). Our occupancy-adjusted model of burrow density estimated a total population of 159,575 breeding pairs (95% CI: 140,710-180,968). Methods differ between the two surveys so the figures are not directly comparable, but we can infer trends by comparing their distributions in 1976 and 2018. Prions remain widespread and appear to have increased their range modestly (Figure 2).

**Figure 2:**
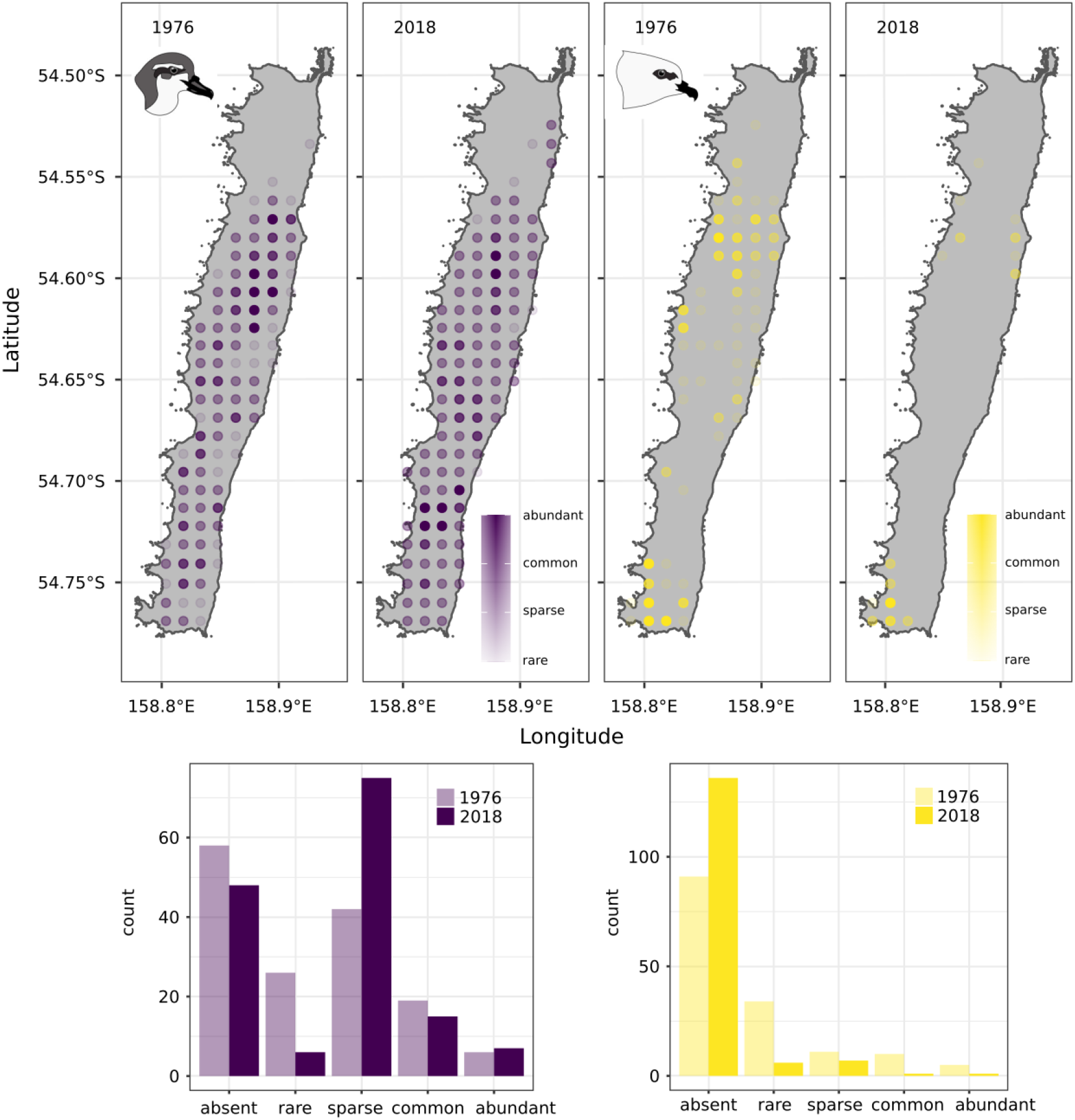
A comparison of Antarctic Prion (left) and White-headed Petrel (right) abundance in 1976 and 2018. Contemporary survey results aggregated by grid cell and scaled using the same percentiles as bins reported by Brothers (1984). The lower plots show change in classified categories.

White-headed Petrels are larger than Antarctic Prions and are present for >9 months of the year from August to June. This increases their vulnerability to cat predation while breeding, resulting in extremely low breeding success when cats were extant (Figure 1; Brothers, 1984; Warham, 1967). Monitoring plots were established in 1993 and revealed a population decline of 14% between 1993 and 1999 prior to cat eradication in 2001 (Brothers and Bone, 2008). Burrow numbers within monitoring plots have since increased above 1993 levels, but comparison of their contemporary distribution with their distribution in the 1970s suggests a marked contraction in their range over the period, which probably took place in the 1980s and 1990s (Figure 2). This is reflected in the reported ratios of Antarctic Prions to White-headed Petrels. Brothers (1984) estimated a prion:petrel ratio of 6:1 in the 1970s. We estimate the ratio is now 13:1.

In the late 1800s Blue Petrels were described as “exceedingly numerous” on the lower coastal slopes of Macquarie Island (Campbell, 1901). By the 1970s they had been extirpated and survived only on offshore rock stacks (Brothers, 1984). Superficially they are similar to Antarctic Prions, breeding in small burrows during the summer months (Bonadonna and Mardon, 2010). So why did prions persist while Blue Petrels perished? In part, it appears to be because Blue Petrels are present for more months in the year re-occupying burrows at the end of the breeding season, and in part because they are restricted at high latitudes to low elevations (Dilley et al., 2017). While there were undoubtedly impacts from rats and cats, Wekas *Galliralus australis*, in particular, are implicated in the complete removal of formerly abundant small petrels from coastal slopes of Macquarie (Brothers and Skira, 1984). Wekas avoided cat predation by occupying dense tussock grasslands around the coast, rarely occurring on the exposed plateau where vegetation is low and sparse. They damaged breeding burrows and predated eggs and chicks (Falla, 1937). The low elevation Blue Petrels succumbed, whereas Antarctic Prions survived in higher elevation refugial habitat on the plateau away from Wekas.

Grey Petrels have large burrow entrances and breed in winter. These two factors make them highly susceptible to cat predation (Barbraud et al., 2009). They were rare though still widespread on Macquarie by 1911 but were extirpated subsequently (Falla, 1937; Schulz et al., 2006).

Species declines in the presence of Wekas and cats, and their survival or extinction were governed by body size, breeding phenology, and the width of their realized niche on the island. Species whose breeding cycles were better suited to intra-annual variation in predation pressure and with broader niches which allowed them to use refugial habitats survived.

### ii) Timing and magnitude of responses to eradications

Since the eradication of cats in 2001 all four species have increased, but their rates and spatial patterns of response have varied. These differences reflect individual responses to changes in rodent and rabbit populations in the early 2000s, and species-specific influences of immigration and philopatry.

Our time series can be split into three stages: stage 1 – pre-cat eradication; stage 2 – post-cat, pre-rodent/rabbit eradication; and stage 3 – post-rodent/rabbit eradication. Following low breeding success and population decline in stage 1 (Brothers and Bone, 2008) White-headed Petrel breeding success increased dramatically to 82% in 1999 when cat numbers fell (Figure 1). Coinciding with cat management Grey Petrels re-established as a breeding species in the 1990s, increasing to a minimum of 59 breeding pairs by 2003 (Schulz et al., 2006). Grey Petrel breeding success rose to 48% in 2004 (Figure 1). Blue Petrel burrows were found on the main island in 1999 (Brothers and Bone, 2008). However, following these early gains at the beginning of stage 2, rabbit numbers increased and the vegetation and soil integrity in breeding petrel colonies on coastal slopes was severely compromised (see Terauds et al., 2014). Breeding success declined again, in White-headed Petrels to a low of 26% in 2010 and in Grey Petrels to 23% in 2007 (Figure 3). Although present, Blue Petrels were not recorded breeding successfully on the main island at this time, likely due to rat predation (Brothers and Bone, 2008). Stage 3 following rabbit and rodent eradication in 2011-2014 has seen breeding success increase to 83% in White-headed Petrels and 74% in Grey Petrels in 2018.

**Figure 3:**
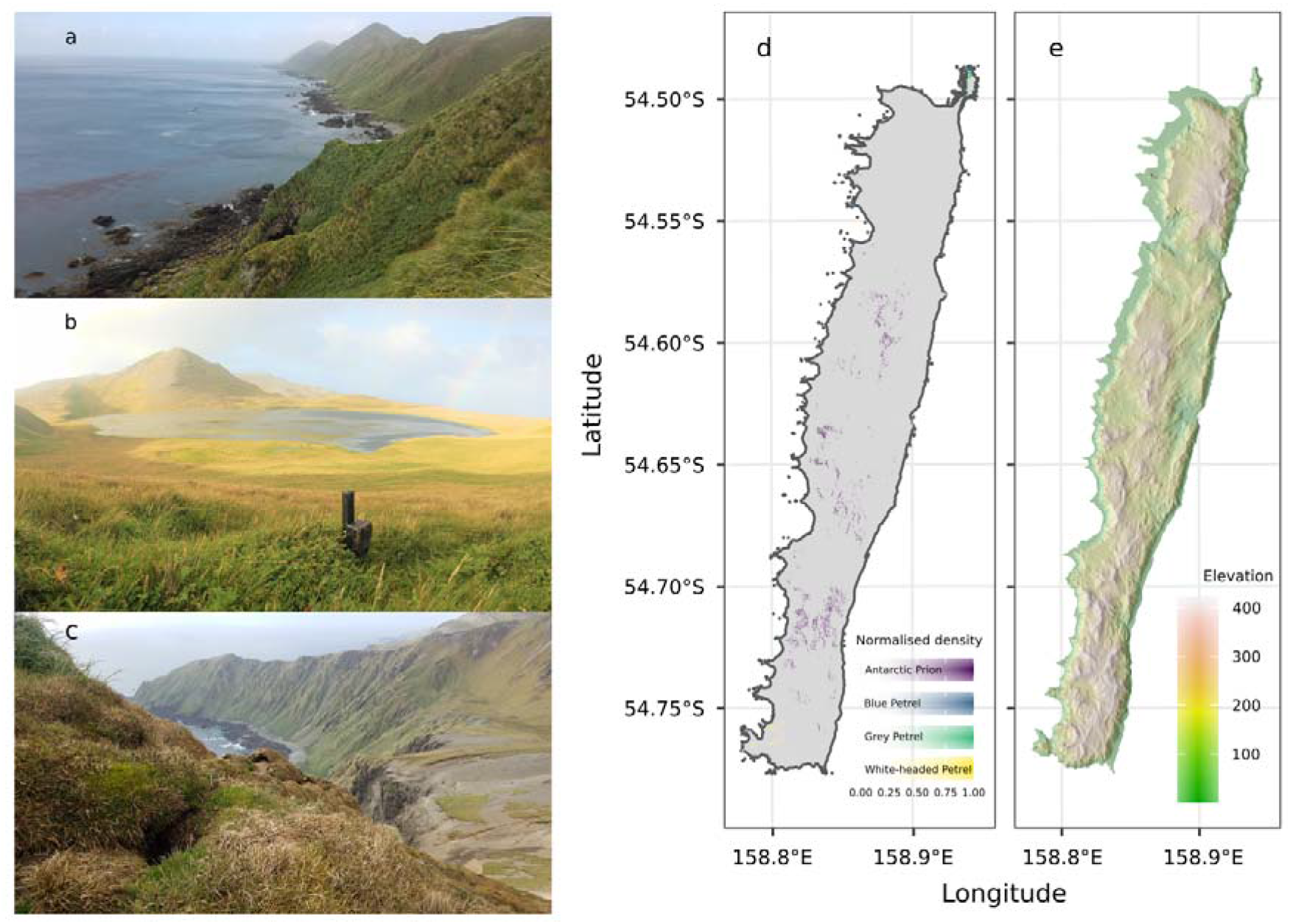
Occupied habitat: a) a mixed Grey and Blue Petrel colony (foreground) with extensive unoccupied coastal slopes (background), b) an Antarctic Prion colony on gentle vegetated slopes on the plateau, c) a White-headed Petrel burrow in a colony on the upper edge of the coastal escarpment, d) combined occupied area for all four species derived from 2018 surveys (Bird et al. submitted), e) a shaded relief map showing the coastal flats, coastal slopes and inland plateau.

Breeding success is a better short-term indicator as it is more responsive to environmental change than burrow numbers because pairs and burrows may be present even when birds are unable to breed successfully. The increase in White-headed Petrel burrow numbers is equivalent to λ = 1.01 (sensu Brooke et al., 2018) since 1993. A smooth of Grey Petrel burrow numbers fluctuates in time with environmental changes, showing a dip during peak rabbit numbers followed by subsequent increases since eradication (Figure 1). Population growth averaged across stages 2 and 3 equates to λ = 1.10 (Table 1). Blue Petrels have not been monitored within plots, but the population has increased rapidly from just 500-600 pairs estimated on offshore rock stacks until 1999 when the first mainland burrows were found, to 5,588 breeding pairs estimated today, a rate of increase of λ = 1.14 (Table 1). Monitoring plots were established for Antarctic Prions in 1988. When smoothed, the relationship between burrow numbers in plots and year becomes linear. Burrows have increased modestly in monitoring plots established in 1988 (Figure 1), from an initial count of 197 burrows to 277 in 2017. Counts have fluctuated since the 1980s with a low of 130 burrows in 1994 and a high of 278 in 2007 (mean 199 ± SD 46), with an overall annual rate of population increase of λ = 1.01. In general prions suffered little from the expansion of the rabbit population in the early 2000s, being able to breed sympatrically (Brothers and Bone, 2008; Raymond et al., 2011).

The two recolonising species, Blue and Grey Petrel, are increasing much more rapidly than the two established species (Figure 1). Petrels are strongly philopatric (Warham, 1996, 1990), but immigration has been shown to play an important role in post-eradication population growth, and may differ markedly between species (Brooke et al., 2018; Buxton, 2014). Rates of Grey and Blue Petrel population increase on Macquarie Island reflect other petrels’ responses to predator removal where immigration is a factor, and exceed those recorded in populations in the absence of immigration (Bonnaud et al., 2009; Brooke et al., 2018). We estimated a constant rate of annual immigration of 20-30 pairs of Grey Petrels since the first burrows were detected in 1993 would have accumulated to the 630 burrows counted in 2018 even without recruitment. In larger populations such low level rates of immigration have a negligible impact on overall population growth rate whereas recruitment and the intrinsic rate of population growth are likely to be more influential (Brooke et al., 2018). Rates of White-headed Petrel and Antarctic Prion population growth support this—both are slightly above intrinsic rates of population growth recorded in closed populations (Bonnaud et al., 2009; Brooke et al., 2018).

Burrow occupancy rates differ between the four species. Occupancy was highest in Antarctic Prions (Table 1) reflecting a higher breeding to non-breeding ratio in the population than for the other species. This is logical for the two recolonising species. As immigration appears to be a major driver of current population growth a large part of the population is likely to be prospecting, unpaired birds, and because the age of first breeding is ~7 in these species (Bird et al., 2020) another large component of the population will be immature. The lower growth rate and larger established White-headed Petrel population suggests immigration is proportionally less in this species. However, high breeding success in recent years may have created a boom of immature prospecting birds, which may result in an increase in population growth rate in the next few years.

### iii) Current distributions and habitat use

The distributions and habitat utilisation of established and recolonising species differ markedly. Distributions are still shaped primarily by the legacy of invasive species impacts. Habitat use reflects legacy and opportunity as shaped by philopatry and immigration.

Antarctic Prions and White-headed Petrels are still confined to refugial habitat. The former to the inland plateau above 200 m, the latter to a few areas of the coastal escarpment at 100-250 m elevation (Figures 3 & 4). Antarctic Prions, which were the least impacted by invasive species, are still widespread with >1 breeding pair predicted to occur currently in 16% of 20 × 20 m pixels across the island. The range of White-headed Petrels has contracted. They are estimated to occur in 0.9% of 20 × 20 m pixels island-wide.

Although they are apparently increasing rapidly, the two recolonising species are still numerically and spatially rare. We estimate the Grey Petrel population to be just 252 pairs today found in only 0.03% of all pixels, mainly on seaward-facing slopes, particularly on the west coast (Figure 1). Blue Petrels have increased to >5,000 breeding pairs but occur in just 0.05% of pixels island-wide with all new colonies on lower coastal slopes around three main focal areas.

There is little overlap in the areas occupied by the four species (Table 2). About one third of pixels occupied by White-headed Petrels also have Antarctic Prions, but only 2% of prion pixels have White-headed Petrels. Approximately 30% and 23% respectively of Grey and Blue Petrel pixels also support the other species.

**Table 2:** Spatial overlap expressed as percentages in the proportion of occupied pixels between species.

|  |  | Occupier |  |  |  |
| --- | --- | --- | --- | --- | --- |
|  |  | Antarctic | Blue | Grey | White-headed |
|  |  | Prion | Petrel | Petrel | Petrel |
| **Owner** | **Antarctic Prion** | 100.0 | 0.0 | <0.01 | 2.3 |
|  | **Blue Petrel** | 0.0 | 100.0 | 22.8 | 0.0 |
|  | **Grey Petrel** | 2.1 | 30.8 | 100.0 | 4.4 |
|  | **White-headed Petrel** | 32.9 | 0.0 | 0.1 | 100.0 |

Despite limited spatial overlap (Table 2) all four species occur in similar vegetation types, primarily short-grass herbfields and tussock grasslands, indicated by the similar NDVI values used by all populations (Figure 4). There is striking overlap in the slopes, elevations, aspects, proximity to ridgelines and topographic wetness utilised by the two recolonising species, and the two established species, but very little overlap between the two groups (Figure 4).

**Figure 4:**
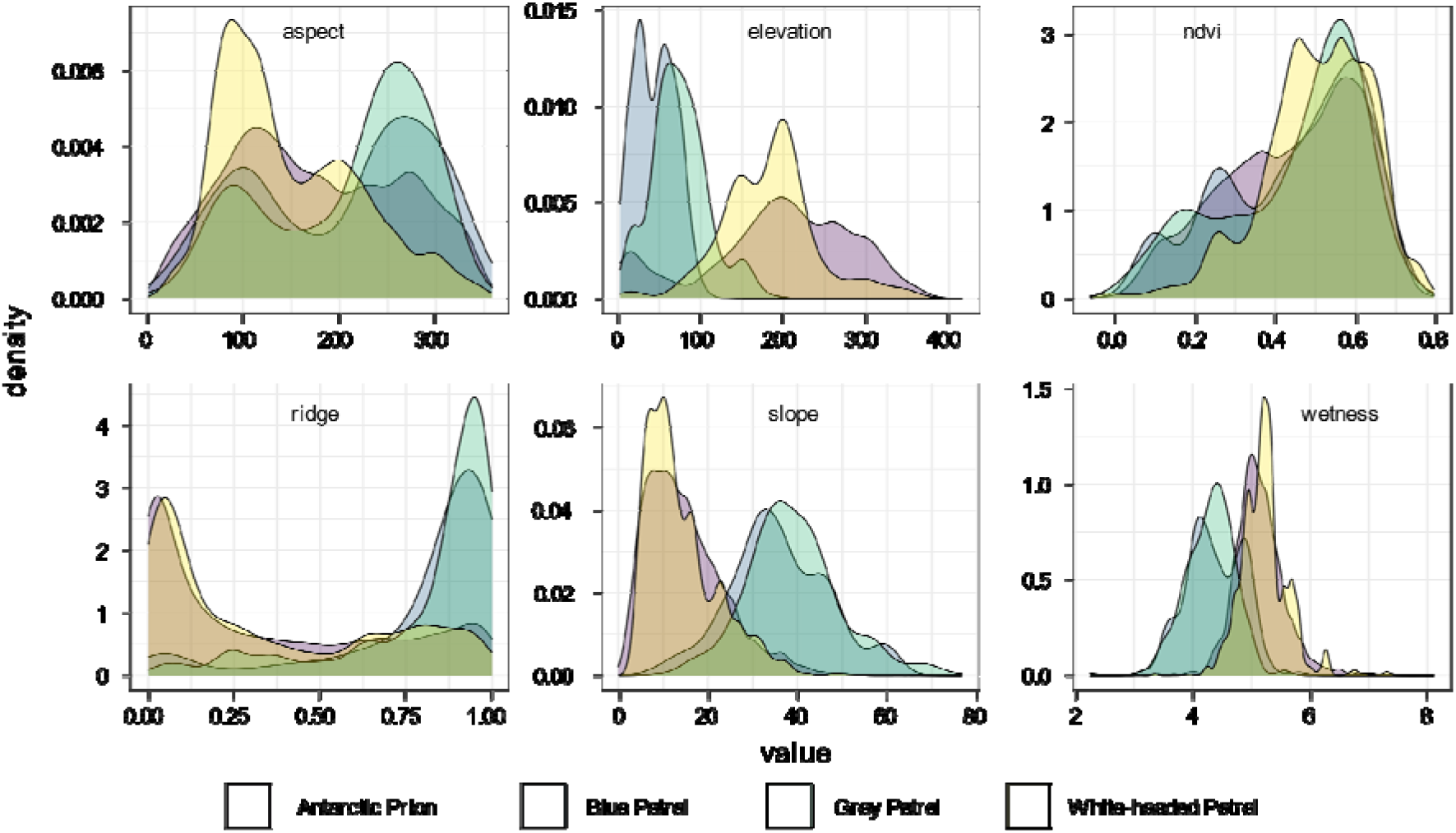
Density plots of proportion of populations in relation to environmental variables. Based upon model-predicted island-wide densities for Antarctic Prions and White-headed Petrels and island-wide search surveys for Blue Petrels and Grey Petrels.

Petrels are non-specific in their breeding habitat requirements, needing only moderately deep, well-drained soils for digging burrows, sufficient vegetation cover to avoid native avian predators, and larger species may favour certain topographies to aid take-off (Schulz et al., 2006; Warham 1990, 1996,). The well-vegetated coastal slopes of Macquarie Island appear to be ideal habitat but are largely devoid of breeding petrels today (Figure 3). The higher elevations, and shallower, wetter slopes away from ridgelines that are used by prions and White-headed Petrels correspond with the island’s plateau. Still apparently confined mainly to refugial areas the niche currently used by both established species may be sub-optimal.

We hypothesise that legacy and opportunity manifest differently for recolonising and established species owing to differential influences of immigration and philopatry. Now that the coastal slopes are free of invasive predators it is to be expected that petrels will recolonise. Indeed, all areas newly occupied by recolonising species are on coastal slopes, reflected in the steeper slopes, lower elevation, lower wetness and closeness to ridgelines of areas used by Blue and Grey Petrels (Figure 4). There were no existing colonies to attract prospecting birds of either species, so their distributions likely reflect their exploitation of prime habitat. It appears to be simply time since eradication that means these slopes are still heavily under-utilised by recolonising species. In contrast, as philopatric colonial nesters, with existing colonies around the upper coastal escarpment and inland plateau, recruiting or immigrating Antarctic Prions and White-headed Petrels are likely to be attracted to existing established sites, even if habitat is sub-optimal (Kildaw et al., 2005). This may result in colony inertia, slowing the rate at which prions and White-headed Petrels recolonise coastal slopes, despite being the most abundant species on the island.

There is further variation at the species level. For example, spatial patterns of recolonization vary markedly between the two recolonising species. Blue Petrels have established at a few focal sites where their colonies are expanding, whereas Grey Petrels have established small new colonies at many sites. Collectively, immigration, breeding success and recruitment, and modes of recolonization appear to be shaping petrel populations on Macquarie Island. Current rates of population growth driven by immigration, and the plasticity of recolonising species to exploit new areas suggests they may close the population gap on established species. Over time we expect spatial overlap to increase to reflect co-occurrence elsewhere (Bonadonna and Mardon, 2010) with the first observations of Antarctic Prions visiting Blue Petrel colonies, and a White-headed Petrel at a Grey Petrel burrow outside their current breeding range recorded in this study (Bird et al. submitted B).

### Implications

Native species’ vulnerability to invasive species impacts is not uniform but can be predicted from their physiology and life histories. Their relative susceptibility foreshadows differential responses to eradication. These differential responses to invasive species and their eradication have implications for impact assessment and forecasting outcomes. In established and recolonising species, the interplay between invasive species legacy and eradication opportunity governs proximate responses.

Immigration is the main influence on recolonising populations, promoting high initial population growth rates and plasticity to colonising optimal habitat in new areas. Philopatry is the main influence on established populations leading to colony inertia which potentially suppresses recovery in sub-optimal habitat.

Relative inertia or plasticity to colonising new areas may increase population sensitivity to emerging threats. For example, although currently Antarctic Prions are the most abundant species on Macquarie, they breed on flatter slopes on the plateau with a higher topographic wetness index so their population may be most susceptible to burrow flooding and breeding failure associated with the increasing frequency of high intensity rainfall events (Brothers and Bone, 2008). Alternatively, small, disparate populations of a recolonising species like Grey Petrel may be more susceptible to predator pits than a more aggregated one like Blue Petrel (Clark et al., 2020).

Invasive species eradication is a highly effective tool for preventing extinctions. By removing the pre-eminent threat from remote island ecosystems which are otherwise relatively free from anthropogenic impacts it also improves conditions for terrestrial, and inter-connected coastal and pelagic ecosystems to develop new stable states. However, there is a danger of complacency. Much of the evidence of species and ecosystem recovery is unpublished or anecdotal. Funding and political support for post-eradication monitoring is limited, but only through quantitative assessments that help to tell the full stories of eradication outcomes will we be able to continue to develop and improve invasive species eradications.

## Methods

Macquarie Island (54° 30’S, 158° 57’ E) is a 12,785 ha UNESCO World Heritage Site and Tasmanian State Nature Reserve. The climate and vegetation are typically sub-Antarctic (Adams, 2009; Pendlebury and Barnes-Keoghan, 2007; Taylor, 1955). Pest Management was implemented by the Tasmania Parks and Wildlife Service in the 1970s-2000s culminating in the joint federal- and state-funded Macquarie Island Pest Eradication Program in 2011-2014 (Copson and Whinam, 2001; Parks and Wildlife Service, 2014; Robinson and Copson, 2014; Springer, 2016). Monitoring plots were established at two sites for Antarctic Prions and two sites for White-headed Petrels in 1988 and 1993 respectively (Brothers and Bone, 2008), with three additional sites added for White-headed Petrels subsequently (Brothers and Bone, 2008; DPIPWE, unpublished data). Grey Petrel colonies have been monitored since 2001 (Schulz et al., 2006; DPIPWE, unpublished data).

We repeated surveys of prion plots in January 2018 and made three visits to White-headed Petrel plots between December 2017 and April 2019, and four visits to Grey Petrel colonies between May and November 2018 to measure breeding success. We established seven monitoring plots for Blue Petrels visited twice in November 2017 and February 2018. Plot visits involved exhaustive searches within the marked boundaries to locate, waypoint and mark all burrows, with repeat searches during revisits used to check type 1 error. All Grey and White-headed Petrel burrows were manually inspected on each visit using a burrowscope or digital camera and the burrowscope and playback were used to determine occupancy in Blue Petrel and Antarctic Prion burrows (see Bird et al. submitted B for full methods and results). In total, monitoring data from 12 Antarctic Prion, 13 White-headed Petrel and 10 Grey Petrel seasons were combined for trend analysis.

We distance-sampled all burrows along transects during a whole-island stratified randomised survey in 2018 from which we derived density surface models of Antarctic Prion and White-headed Petrel abundance, and completed whole-island targeted search surveys for Grey Petrels and Blue Petrels in May-October and August-October respectively (see Bird et al. submitted A for full survey methods and results).

We fitted a generalized additive model to burrow counts and estimated breeding success across all years with species as a factor and site as a random effect (Pedersen et al., 2019). We calculated population change (λ – lambda) as the *x*^*th*^ root of the proportional population change between previous surveys/census and our 2018 survey and census of Blue and Grey Petrels (Brooke et al., 2018). For Antarctic Prions and White-headed Petrels λ was calculated from the exponent of the species-specific terms predicted by the model on the link scale, assuming a constant gradient throughout the period. Methodological differences between surveys in 1976 and 2018 precluded direct comparison of Antarctic Prion and White-headed Petrel population estimates, but to compare distributions we re-scaled modelled densities across the island on the same grid used by Brothers (1984). We binned aggregated abundance into five categories: abundant, common, sparse, rare or absent defining the bins using the same percentiles (Brothers, 1984).

All analyses were performed, and plots prepared in R versions 4.0.1 with associated packages mgcv, tidyverse, sf and raster (Hijmans, 2020; Pebesma, 2018; R Core Team, 2020; Wickham et al., 2019; Wood, 2017).

## Authors’ contributions

JB and JS conceived the ideas, JB designed methodology, collected and analysed the data, and led the writing of the manuscript. All authors contributed critically to data interpretation, drafts and gave final approval for publication.

## Acknowledgements

The authors thank Noel Carmichael and Tasmania Parks and Wildlife Service for their support facilitating this project. Thanks to Toby Travers for suggestions for data manipulation and analysis, and to past Wildlife Rangers from Macquarie Island for their help interpreting data, in particular Helen Achurch, Julie McInnes, Marcus Salton and Penny Pascoe. This study was supported by funding from the Australian Government’s National Environmental Science Program through the Threatened Species Recovery Hub, and the Australian Antarctic Science program (AAS 4305). JB was supported by a Research Training Program scholarship, an Antarctic Science International Bursary, National Environmental Science Programme Threatened Species Recovery Hub Research Support and a BirdLife Australia Stuart Leslie Bird Research Award. All methods were approved by the University of Queensland Native/Exotic Wildlife and Marine Animals (NEWMA) animal ethics committees (AE29713), the Macquarie Island Research Advisory Group and the Department of Primary Industries, Parks, Water and Environment (TFA 17305). Access to Macquarie Island was granted by the Tasmania Parks and Wildlife Service (Access Authority No. 17-18 5).

## Data Sharing

Raw data and full code for the analysis are available from the Australian Antarctic Data Centre.

